# Tuning a Genetic Circuit with Double Negative Feedforward Loops to Approximate Square Waves

**DOI:** 10.1101/2025.11.13.688370

**Authors:** Jiayi Wu, Charles Johnson, Enoch Yeung

**Affiliations:** Molecular, Cellular, and Developmental Biology, University of California, Santa Barbara, Santa Barbara, California, 93117, United States; Department of Mechanical Engineering,University of California, Santa Barbara, Santa Barbara, California, 93117, United States

## Abstract

Precise temporal control of gene expression is important for programming cellular functions. Existing circuits, however, often couple amplitude and duration. This prevents independent tuning of key dynamic features. We therefore set out to design a genetic system capable of generating square-wave outputs with externally programmable geometry. To achieve this, we constructed a layered regulatory design that separates production and removal into two independently controlled modules. An activating input turns expression on and builds the reporter. A repressing input stops synthesis and removes existing protein. Together, these actions confine the active window and produce square-wave outputs or pulses on demand. Waveform geometry depends on input dose and timing. Analysis of limiting cases reveals that strong repression compresses dynamic range, and finite degradation capacity restricts decay rate. Enhancing removal capacity alleviates this bottleneck and sharpens the off-transition. Together, this work establishes a general framework for programming gene-expression dynamics with independently tunable amplitude and duration.

## INTRODUCTION

Gene circuits have enabled programming of cellular functions to perform sensing, logic processing, and dynamic responses.^1^ However, achieving predictable temporal control over gene expression remains a central challenge^2^.^3^ Inducible promoters responsive to arabinose, ^4^ tetracyclines^5^,^6^ or IPTG, once triggered, maintain transcription until the inducer is removed or diluted out. Autonomously timed circuits such as oscillators^7 8^ have instantaneous time courses set by internal feedback rather than by user inputs at runtime. Since constitutive systems and oscillators often intrinsically couple key output features through a single input, this coupling limits our ability to independently tune the amplitude and duration of genetic signals. However, this capability is critical for both natural systems and industrial applications. For example, in eukaryotic cells the NF*κ*B pathway regulates responses to pathogens and stress; brief high-amplitude activation promotes pro-inflammatory targets, whereas lower-amplitude, prolonged activity favors cell survival or tolerance.^9^ Likewise, precise control over the duration and amplitude control of effector output in therapeutic drug delivery can balance efficacy and toxicity while limiting drug resistance^10^.^11^

We therefore aimed to engineer a genetic circuit that can have a programmable square-waveform output characterized by a steep climb to a stable plateau, a steady-phase state, and a rapid decline that allows for a bounded window of activity to exist. Specifically, we not only aimed to synthetically generate this geometry but also to provide independent, external control over its key parameters: amplitude and duration.

In order to realize this goal, we constructed a system based on layered negative regulation. Control was divided into two functionally independent arms: one inducible activation element for ramp-up and two inducible repression elements for ramp-down. Production governs the ramp-up phase and the plateau level. Cooperative, ^12^ inducible activation produces a steep rise and a tunable steady-state plateau. Meanwhile, the reduction of production and the removal of existing proteins govern the duration and rate of ramp-down. A rapid, timed ramp-down is obtained by layering transcriptional inhibition, which halts new production, with a controllable proteolysis module that selectively degrades the target protein^13^.^14^ To decouple these two phases of expression, we applied separate external inputs to control rampup and ramp-down activity. Dual-input control enables a controllable, square-wave-like circuit with tunable amplitude and duration.

We assessed the system through quantitative measures that characterize waveform quality with respect to amplitude, plateau duration, and rise and fall rates. Two critical constraints limit square-wave fidelity. First, overly strong transcriptional inhibition can suppress promoter activity and dampen the amplitude of the output. Second, finite protease capacity leads to saturation at high substrate loads, which restricts the degradation rate and consequently slows the ramp-down transition.^15^ These results indicate that physiological limits of cells can potentially constrain square-waveform characteristics, particularly the steepness of the ramp-down, and the amplitude tunability of the output.

Together, this work introduces a gene circuit for programming gene-expression dynamics through layered negative regulation. By separating production and removal into independently tunable modules, we create a circuit with controllable amplitude and duration that produces square-wave outputs and, on demand, pulse-like behaviors.

## RESULTS

### Layered Transcriptional and Proteolytic Repression Generates a Square Waveform Output

Generating defined temporal programs of gene expression is a central goal in synthetic biology. However, many designs yield broad, poorly defined windows because decay is slow^16 2^. In addition, the amplitude and duration cannot be tuned independently. To bridge the gaps in existing gene circuits, we aimed to generate a gene circuit in which, by applying inputs to the system, the reporter undergoes activation and deactivation in cells and ultimately generates a square waveform output. Moreover, the timing of input addition and the concentration of the input can shape the amplitude and duration of the output (Figure 1A).

**Figure 1.**
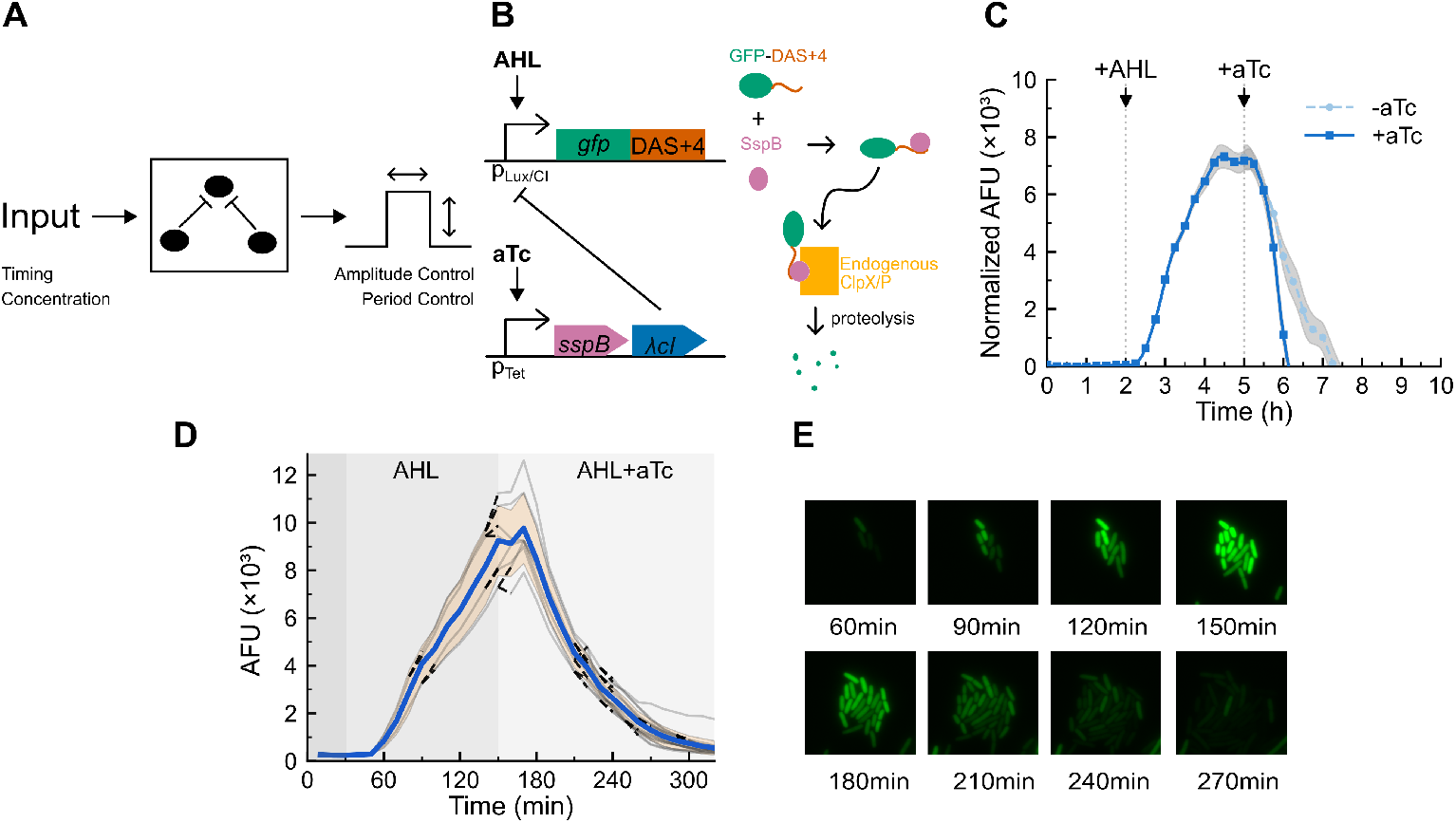
Layered negative regulation produces a square-wave gene-expression output. **(a)** Design schematic showing that input timing and concentration set waveform amplitude and duration. **(b)** Schematic of the two-plasmid system. Left: reporter plasmid and negative-regulator plasmid. GFP fused to a DAS+4 degron is driven by the Lux/*λ*CI promoter on a ColE1 backbone. SspB and *λ*CI are expressed from the Tet promoter on a p15A backbone. Right: SspB adaptor delivers GFP–DAS+4 to endogenous ClpXP for degron-specific proteolysis. **(c)** Plate-reader time courses with activating and repressing inducers. GFP/OD_600_ trajectories from cultures induced with AHL (35 nM at 2 h) and, where indicated, aTc (100 ng mL^−1^ at 5 h). Data are background-subtracted and OD-normalized; curves show mean ± SD (*n* = 4), with gray bands denoting SD. Vertical dotted lines mark inducer addition times; colors correspond to +aTc and –aTc conditions. **(d)** Single-cell fluorescence trajectories. Time-lapse degron-tagged GFP expression curves for individual cells under continuous flow of MOPS medium containing 35 nM AHL followed by 35 nM AHL + 100 ng mL^−1^ aTc. Each line represents one cell. Solid central lines denote the mean across cell lineages within the population, and the shaded region indicates ±1 SD. **(e)** Representative single-cell fluorescence micrographs at the indicated times after induction.

To meet this design objective, we proposed a gene circuit that layers transcriptional repression with targeted proteolysis (Figure 1B). In particular, two convergent negative controls act on the same output to ensure that both transcript production and protein accumulation are actively arrested. Our output signal for the circuit is a GFP protein, regulated by three elements: 1) *λ*CI transcription factor, 2) SspB-regulated degradation, 3) inducible promoters to regulate expression of SspB and *λ*CI. At the transcriptional level, *λ*CI protein dimers halt transcription by binding operator sites on a *λ*CI-Lux hybrid promoter. We tag GFP with a Das+4 degradation tag (DAS+4 is a variant of the ssrA tag). The Das+4 degradation tag does not enable direct binding of the ClpXP protease, but requires SspB adaptor to first bind the GFP-Das+4 protein for delivery to ClpXP-mediated degradation.^14^ Lastly, we use TetR and it’s cognate inducer anhydro-tetracycline to regulate expression of *λ*CI and SspB adaptor protein These combinatorial, transcriptional, and post-translational controls ensure a rapid ramp-down of GFP expression, required for approximating square waves.

We implemented this design as a two-plasmid system in E. coli (Figure 1B). The reporter plasmid (ColE1 origin) contained an AHL-inducible Lux/*λ*CI promoter driving expression of GFP fused to a DAS+4 degron. The regulator plasmid (p15A origin) contains an aTc-inducible Tet promoter driving a polycistronic mRNA encoding the SspB adaptor protein followed by the *λ*CI repressor. In this design, addition of activating inducer (AHL) induces GFP expression. Subsequent addition of repressing inducer (aTc) induces the co-expression of SspB and *λ*CI. SspB delivers the GFP-DAS+4 reporter to the ClpXP protease for degradation, while *λ*CI binds to the Lux/*λ*CI promoter to halt further transcription. These two plasmids were then co-transformed into the *E. coli* strain MG1655ΔSspB, since wildtype *E. coli* strains have endogenous SspB expression.

Having constructed the reporter–regulatory system, we next tested its temporal behavior at the population and single-cell level to determine whether the layered repression design produces a square-wave output. In bulk culture, fluorescence increased to a stable plateau and then returned to baseline when the repression module was induced(Figure 1C). Trapezoid pulse fits were used to extract rise time, plateau duration, plateau height and drop time (see Methods).

We found that we could use aTc induction to reduce the drop time of our circuit’s output by threefold. By contrast, the rise time, plateau height, plateau duration were comparable with induction of aTc versus without. This capacity to independently tune the drop time from the other parameters of the wave output facilitates the use of this circuit in controlled settings. Specifically, the induced group’s drop time lasted 0.67 h, whereas that of the group without induction lasted 2.0 h. This represents a threefold acceleration in the ramp-down phase with layered repression activated, which tightens the active window and makes the waveform more square. Consistent with the population results, single-cell measurements also confirmed that this behavior is not a population-averaging artifact (Figure 1D-1E). Individual trajectories followed the same pattern, with coordinated activation and a return to baseline despite variation in peak amplitude. Together, these results show that combined transcriptional repression and proteolysis produce a square waveform output across scales with a well-defined ramp-down transition.

### Inducer Concentration and Timing Allows Independent Tuning of Square-wave Amplitude and Duration

After verifying that the circuit produces a square waveform output (Figure 1), we inquired whether the two principal geometric properties of the circuit, amplitude and period, are tunable independently. We reasoned that the steady-state level of activation would depend on transcriptional activation strength and duration of the activity window would depend on the timing of repression. To test this, we independently varied the concentration of the activating inducer (AHL) (Figure 2A), and the delay (*τ*) between the addition of AHL and the addition of the repressing inducer (aTc) (Figure 2C). Recall that the activating inducer, AHL, sets the activty of the Lux/*λ*CI promoter, and the repressing inducer, aTc, controls the activation time of the layered negative regulators.

**Figure 2.**
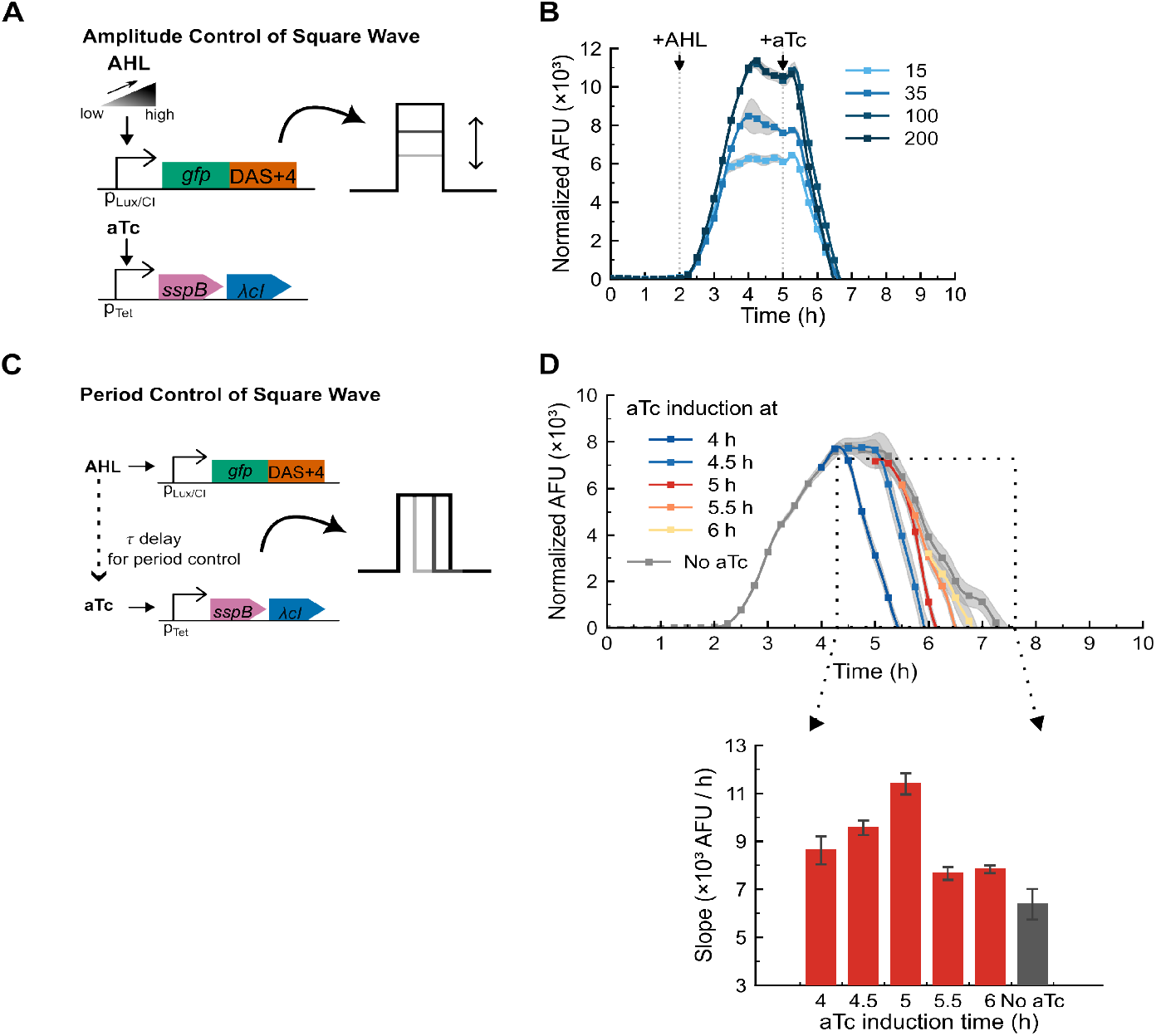
Inducer dose and timing independently tune amplitude and duration of the square-wave output. **(a)** Schematic of amplitude control by AHL dose. **(b)** Plate-reader time courses across AHL doses. GFP/OD_600_ trajectories from cultures induced with AHL at 15, 35, 100, or 200 nM (2 h) and aTc (100 ng mL^−1^ at 5 h). Data are background-subtracted and OD-normalized; curves show mean ± SD (*n* = 4), with gray bands denoting SD. Vertical dotted lines mark inducer addition times; colors correspond to AHL doses. **(c)** Schematic of period control by aTc timing (*τ* delay). **(d)** Plate-reader time courses at varying aTc induction times with slope summary. GFP/OD_600_ trajectories with AHL fixed at 35 nM (2 h) and aTc added at 4, 4.5, 5, 5.5, or 6 h; “No aTc” control included. Data are background-subtracted and OD-normalized; curves show mean ± SD (*n* = 4), with gray bands denoting SD. Lower panel: ramp-off slopes computed by linear fits to the decline segment within the dotted window; bars show mean ± SD (*n* = 4).

We first investigated whether AHL concentration regulates amplitude without affecting the waveform duration. Increasing AHL from 15 to 200 nM raised the fluorescence peak progressively and the response saturated between 100 and 200 nM (Figure 2B). Peak fluorescence increased from approximately 6.5 *×* 10^3^ at 15 nM AHL to 1.1 *×* 10^4^ AFU at 100 nM, corresponding to an approximately 1.7-fold range in tunable amplitude. When we fit trapezoid pulses to these waves (see Methods for details), the parameters of our best fit found similar rampup times, ramp-down times and plateau durations across doses, indicating that promoter activity drives the change in response amplitude and that changing the amplitude does not alter the duration or the ramp-up or ramp-down kinetics. These data indicate that amplitude is continuously tunable with AHL concentration, with saturation suggesting full occupancy of the Lux/*λ*CI promoter.

We next tuned the duration of the plateau and observed that we could independently increase the plateau duration of the wave adding aTc after a delay. Holding the AHL concentration and the addition time constant, we added aTc at 30-minute intervals from 4 h to 6 h to each group of biological replicates (Figure 2D). Earlier induction at 4 h produced a 0.31 h plateau duration, whereas later induction extended the plateau before decay, specifically, induction at 4.5 h and 5 h had 0.89 h and 1.35 h plateau durations, respectively. In addition to altering the plateau duration, there was also a correlation between the ramp-down rate and the timing of aTc addition. Additions between 4 and 5 h yielded progressively steeper ramp-down rates, with the steepest around 5 h and decreasing again after 5 h. Thus, the timing of aTc addition primarily determines the plateau duration and the ramp-down kinetics. We therefore conclude that the two principal geometric properties of the circuit, amplitude and period, are independently tunable.

### Programmable Pulse Generation through Early Repression

Having established independent control of square-wave amplitude and period (Figure 2), we next asked whether the same circuit could also produce a single pulse, similar to other pulse generating circuits previously published.^12^ We hypothesized that inducing the negative regulators before the plateau forms could generate a pulse. We also asked whether pulse amplitude and timing would remain programmable (Figure 3).

**Figure 3.**
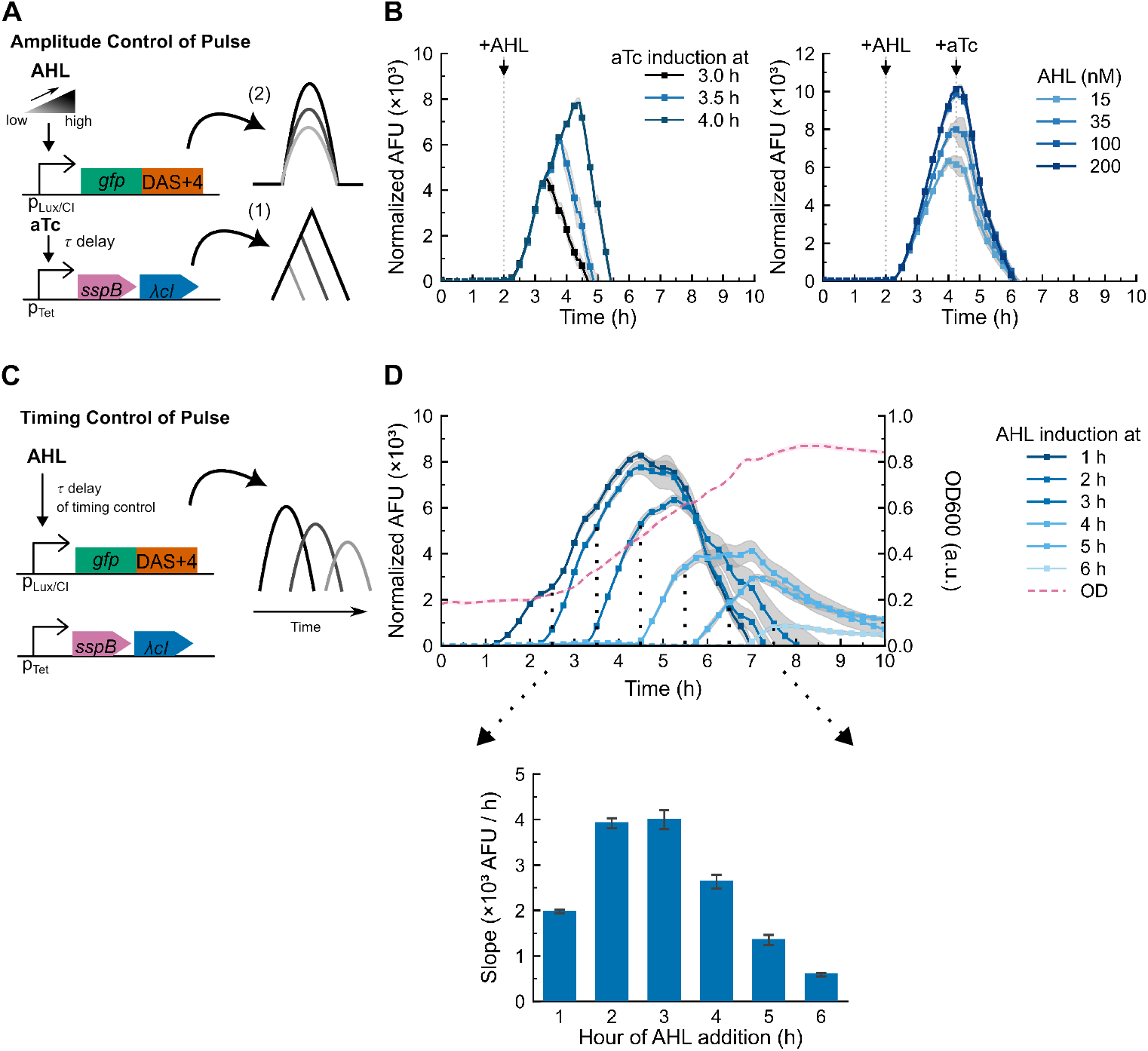
Pulse generation and control. **(a)** Schematic of pulse generation by early repression and amplitude control by AHL dose. **(b)** Plate-reader time courses for pulse experiments. Left: GFP/OD_600_ trajectories from cultures induced with AHL (35 nM at 2 h) and aTc (100 ng mL^−1^) added at 3.0, 3.5, or 4.0 h. Right: GFP/OD_600_ trajectories from cultures induced with AHL at 15, 35, 100, or 200 nM (2 h) and aTc (100 ng mL^−1^ at 4.0 h). Data are background-subtracted and OD-normalized; curves show mean ± SD (*n* = 4), with gray bands denoting SD. Vertical dotted lines mark inducer addition times; legends indicate aTc timing (left) or AHL dose (right). **(c)** Schematic of pulse timing control by AHL onset (*τ* delay). **(d)** Plate-reader time courses at varying AHL induction times with ramp-up slope summary. GFP/OD_600_ trajectories from cultures induced with AHL at 1, 2, 3, 4, 5, or 6 h in the absence of aTc; OD (dashed) shown for reference. Data are background-subtracted and OD-normalized; curves show mean ± SD (*n* = 4), with gray bands denoting SD. Lower panel: ramp-up slopes computed by linear fits over the first 1.5 h after AHL addition; bars show mean ± SD (*n* = 4).

To test whether the same circuit could generate a pulse-like output, we activated the negative regulators during the rising phase of the reporter signal and found that the square wave collapsed into a pulse and the amplitude remained tunable. With the AHL addition time and concentration kept constant, we activated the negative regulators at 3.0, 3.5, or 4.0 h. Each condition produced a transient pulse with a rapid decline(Figure 3A-3B). Using trapezoid fits^17^,^18^ the steady-state windows for the 3.0 h and 3.5 h groups were approximately 0.05 h and 0.31 h for the 4.0 h group, which confirms pulse behavior. With a fixed aTc time, raising the AHL concentration (15, 35, 100, 200 nM) increased the pulse amplitude, while the length of the steady-state window remained the same (Figure 3B). Together, early repression converts the square wave into a pulse, and the amplitude remains tunable by the AHL concentration.

Pulse amplitude also depends on how much reporter has accumulated before repression begins, since the repressive inducer is added early to prevent it from entering the maximum steady state and thereby generating a pulse. We next aimed to quantify how activation timing shapes the rising phase. We found that the timing of activating inducer (AHL) addition determines not only the phase of activation but also the production rate and the potential maximum amplitude attainable before repression. We varied the AHL addition time from 1 to 6 h in the absence of aTc and quantified the ramp-up rate for each condition over the first 1.5 h after induction (Figure 3C-3D). Later AHL addition shifted the response proportionally in time and altered the ramp-up rate. Induction at 1 h yielded a ramp-up slope of approximately 2 × 10^3^ AFU per 1.5 h, whereas induction at 2 h and 3 h roughly doubled this value(Figure 3D). Beyond 3 h, the slope progressively declined with later induction (4–6 h). Moreover, earlier addition of the activating inducer allowed a longer GFP production window and thus yielded a higher maximum amplitude(Figure 3D). These results therefore show how activation timing establishes the kinetic ceiling that limits the achievable pulse amplitude under early repression.

Together, these results show that two inputs define pulse geometry. The repressing input truncates expression to set the pulse, while the activating input sets amplitude and phase via its dose and onset time.

### Dissecting Transcriptional Repression and Proteolysis Reveals Complementary Limits

After the complete gene circuit generated a square-wave output, we wondered how SspB adaptor protein–mediated targeted proteolysis and *λ*CI-mediated transcriptional repression contribute to the ramp-down rate, respectively. To investigate this question, we constructed two control strains. One strain contained only the *λ*CI coding sequence under the control of the Tet promoter on the regulatory plasmid(Figure 4A), while the other contained only the SspB coding sequence under the same conditions (Figure 4C). This isolation of each negative regulator’s effect on waveform shaping provides insight into their respective contributions to square waveform generation.

**Figure 4.**
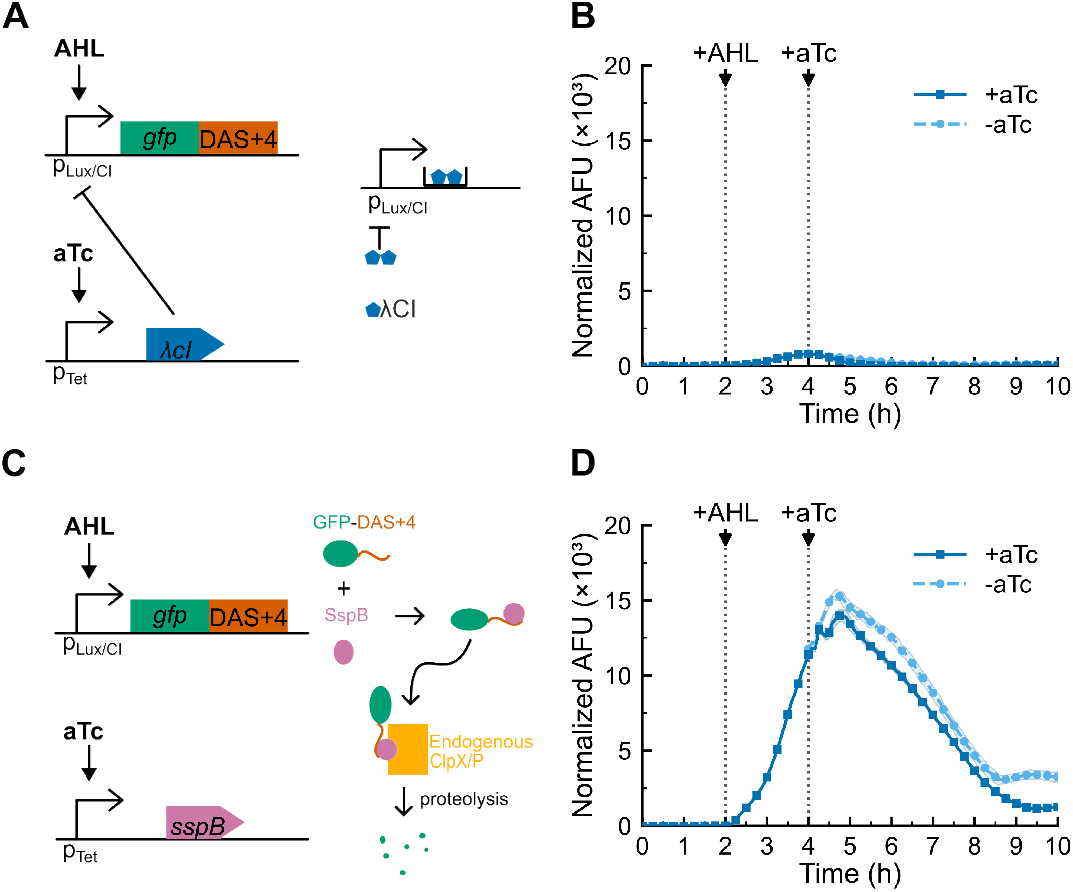
Single-regulator controls reveal distinct limitations. **(a)** Schematic of *λ*CI-only control strain. The regulator plasmid encodes *λ*CI under control of the Tet promoter (*P*_tet_); SspB is absent. **(b)** Plate-reader time courses for *λ*CI-only strain. GFP/OD_600_ trajectories from cultures induced with AHL (35 nM at 2 h) and, where indicated, aTc (100 ng mL^−1^ at 4 h). Data are background-subtracted and OD-normalized; curves show mean ± SD (*n* = 4), with gray bands denoting SD. Vertical dotted lines mark inducer addition times; colors correspond to ± aTc conditions. **(c)** Schematic of SspB-only control strain. The regulator plasmid encodes SspB under control of the Tet promoter (*P*_tet_); *λ*CI is absent. **(d)** Plate-reader time courses for SspB-only strain. GFP/OD_600_ trajectories from cultures induced with AHL (35 nM at 2 h) and, where indicated, aTc (100 ng mL^−1^ at 4 h). Data are background-subtracted and OD-normalized; curves show mean ± SD (*n* = 4), with gray bands denoting SD. Vertical dotted lines mark inducer addition times; colors correspond to ±aTc conditions.

Contrary to our initial hypothesis that two negative regulators would each partially reproduce the ramp-down kinetics, the two modules instead displayed opposing behaviors. In the strain that only has *λ*CI as a negative regulator, the amplitude of the output signal was suppressed significantly even without the addition of the repressive inducer (aTc), suggesting strong basal repression that abolishes the dynamic range (Figure 4B). By contrast, although the strain that only expresses SspB had higher amplitude, induction of SspB only settled the fluorescence trajectory at a lower plateau and then had a comparable ramp-down slope to that without SspB induction (Figure 4D). Thus, neither control recapitulates the behavior of the complete circuit and each exposes a distinct limitation which is over-repression by *λ*CI and minimal contribution of SspB to the complete circuit’s ramp-down rate under endogenous ClpXP capacity.

### ClpXP Supplementation Relieves the Protease Bottleneck to Accelerate Rampdown

Given that SspB-mediated targeted proteolysis had a limited effect on accelerating the rampdown rate, we hypothesized that endogenous ClpXP may be saturated by high-copy-number substrates, thereby capping protease capacity. Therefore, we constructed a new regulatory plasmid that places SspB and ClpXP under different inducible promoters to test whether external supplementation of ClpXP can increase protease capacity (Figure 5A). The results showed that supplementing either component alone was insufficient to accelerate the ramp-down rate (Figure 5B). Specifically, inducing SspB or ClpXP alone caused an earlier decline in the trajectories after the plateau, but the ramp-down rate remained comparable to that of the no-inducer control. Notably, when SspB and ClpXP were co-induced, the rampdown rate increased significantly and the signal returned to baseline earliest. Taken together, these results indicate that protease capacity limits the ramp-down rate and that supplementing ClpXP relieves this bottleneck.

**Figure 5.**
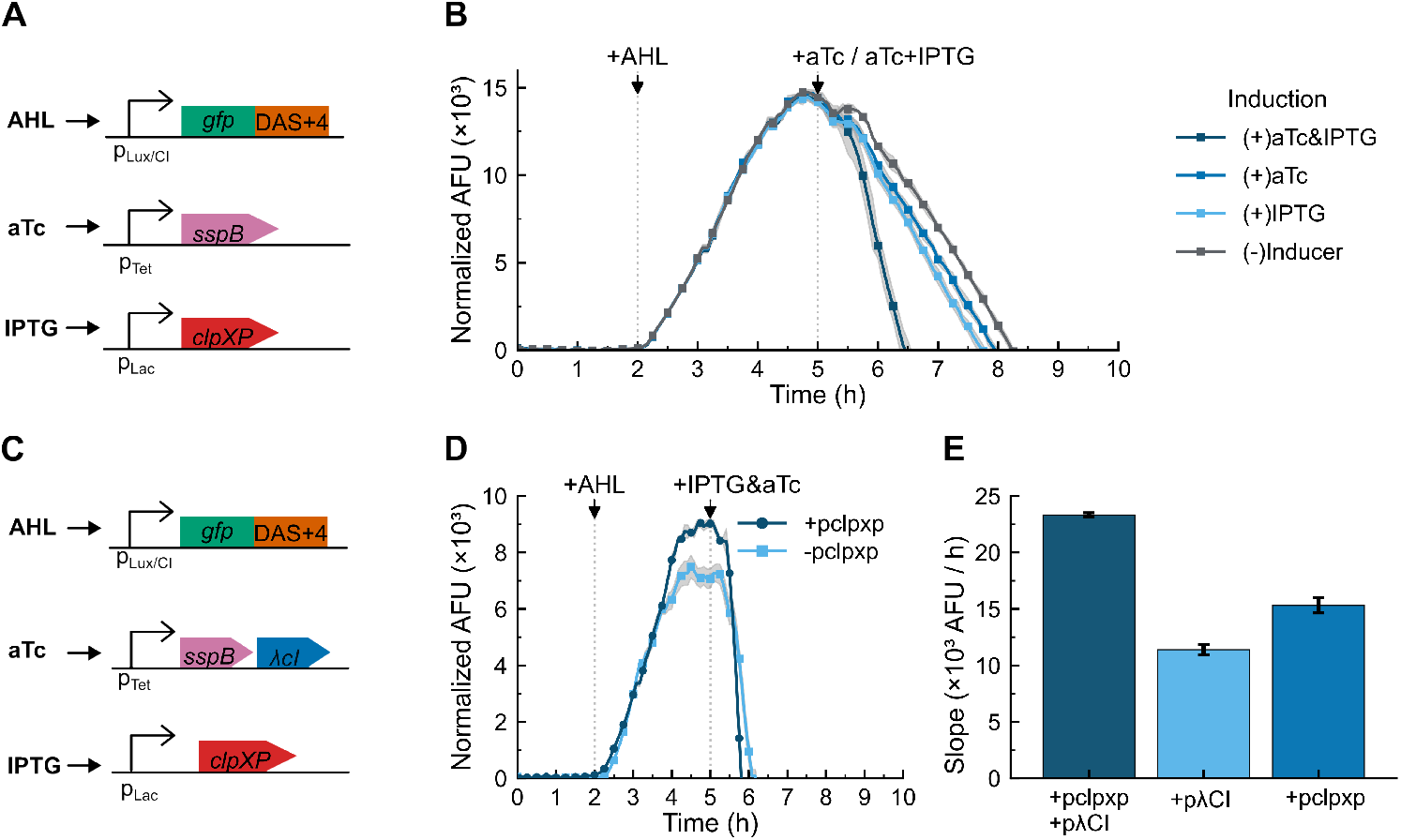
Increasing protease capacity accelerates the ramp-off rate. **(a)** Schematic of SspB–ClpXP co-induction design. The reporter (GFP–DAS+4) is expressed from the AHL-inducible Lux/*λ*CI promoter. SspB and ClpXP are placed under the Tet and Lac promoters, respectively, allowing independent induction by aTc and IPTG. **(b)** Platereader time courses for ClpXP supplementation. GFP/OD_600_ trajectories from cultures induced with AHL (35 nM at 2 h) and, where indicated, aTc (100 ng mL^−1^ at 5 h) and/or IPTG (1 mM at 5 h). Data are background-subtracted and OD-normalized; curves show mean ± SD (*n* = 4), with gray bands denoting SD. Vertical dotted lines mark inducer addition times; colors correspond to induction combinations. **(c)** Schematic of complete layered circuit with ClpXP supplementation. The reporter, SspB–*λ*CI, and ClpXP modules are under Lux/*λ*CI, Tet, and Lac promoters, respectively. **(d)** Plate-reader time courses comparing ± *p*ClpXP conditions. GFP/OD_600_ trajectories from cultures induced with AHL (35 nM at 2 h) and aTc + IPTG (100 ng mL^−1^ and 1 mM at 5 h). Data are background-subtracted and OD-normalized; curves show mean ± SD (*n* = 4), with gray bands denoting SD. Vertical dotted lines mark inducer addition times; colors correspond to the presence of ClpXP on the regulatory plasmid. **(e)** Bar plot of maximum ramp-off rate. Bars show mean ± SD (*n* = 4) of the maximum ramp-off rate calculated by linear regression. All groups carry SspB; colors correspond to the presence of *λ*CI, ClpXP, or both on the regulatory plasmid.

Building on this result, we further accelerate ramp-down rate using targeted genetic design. We added ClpXP to the complete circuit that contains both SspB and *λ*CI (Figure 5C). Our hypothesis was that increasing protease capacity could further accelerate the ramp-down rate. We found that co-induction of SspB, ClpXP, and *λ*CI increased the ramp-down rate by approximately twofold relative to the configuration lacking plasmid-borne ClpXP, and by approximately 1.5-fold relative to the configuration lacking *λ*CI (Figure 5E). In other words, simultaneous induction of transcriptional repression and enhanced protease capacity for SspB-mediated, degron-specific proteolysis maximizes the ramp-down kinetics.

## DISCUSSION

Our findings show that a layered negativecontrol design combining transcriptional repression and targeted proteolysis enables externally programmable waveform geometry. In particular, transcriptional repression halts further protein accumulation, while targeted degradation accelerate the removal of existing proteins; together, these effects result in a high rampdown rate, a defining square-wave characteristic that enables tight-window control of expression. Building on this design, we separate production from removal to achieve orthogonal control of amplitude and duration. This separation converts a typically coupled one-input system into a two-parameter control system that allows independent programming of timing and amplitude, thereby realizing square-wave and pulse outputs.

The construct spans a continuum of temporal profiles. Early repression generates a pulse, with amplitude determined by activation strength and the short pre-repression accumulation period. Delaying repression until after the rise yields a square-wave output; the amplitude remains set by activation, whereas the duration is set by repression timing.

However, two mechanistic limits affect waveform fidelity. First, strong repressor activity suppresses the amplitude of the output and diminishes the dynamic range. When *λ*CI was placed directly under the Tet promoter, basal repression was sufficient to suppress the amplitude even without the addition of an inducer. The over-repression did not appear when *λ*CI followed SspB in a polycistron. Therefore, we hypothesize that translational coupling attenuates *λ*CI’s basal production, where ribosome re-initiation downstream of SspB reduces effective *λ*CI translation in the absence of induction, thereby lowering leak and preserving dynamic range. Second, finite protease capacity constrains the off-transition. ClpXP saturation by high-copy substrates is hypothesized to cause the failure of activation of the SspB adaptor protein to accelerate. The relief of this bottleneck by supplementing ClpXP from a plasmid indicates that protease capacity, not degradation tag recognition by SspB alone, limits the ramp-off rate.

Taken together, these findings suggest practical design rules. First, controlling production and removal with orthogonal inputs provides independent control over amplitude and duration. Second, stopping transcription and actively removing existing protein bounds the activity window and steepens the off-transition. Third, regulator stoichiometry can be tuned at the level of genetic architecture; arranging *λ*CI downstream in a polycistronic transcript can reduce basal repressor accumulation and thereby preserve dynamic range without altering the promoter’s operator affinity. Finally, physiological capacity can be limiting; in our system, endogenous ClpXP constrained the off-transition rather than SspB expression.

Although these design rules resolve key mechanistic constraints such as over-repression and limited proteolysis capacity, several broader limitations remain. The circuit’s behavior depends on the host context because protease abundance, cellular burden, and competition from endogenous substrates can shift the effective clearance capacity. In addition, substrate specificity limits generality, since our reporter is readily degraded by ClpXP whereas other proteins may require alternative proteases or tag modifications to achieve comparable clearance efficiency.

The design principles demonstrated here can be extended to systems that require precise temporal control, including therapeutic delivery, staged metabolic production, and developmental programs with time-gated gene expression. Because the production and removal modules operate independently, this architecture can be adapted to other repressors, proteases, or reporters while preserving the same control logic. More broadly, it provides a compact framework for constructing synthetic circuits with programmable activity windows and sharp shutoff dynamics.

## METHODS

### Plasmid Assembly & Strain Development

All plasmids used in this study were constructed by Gibson assembly. DNA fragments were synthesized by Integrated DNA Technologies (IDT) and assembled into two plasmid backbones originally provided by Glen E. Cronan and Andrei Kuzminov (University of Illinois at Urbana–Champaign): a p15A-origin plasmid carrying P_tet_-driven *sspB* and a ColE1-origin plasmid carrying GFP fused to a DAS+4 degron.

Plasmids were first transformed into *Escherichia coli* DH5*α* for propagation and sequence verification by whole-plasmid sequencing (Eurofins Genomics). Verified constructs were subsequently transformed into *E. coli* K-12 MG1655 Δ*sspB* (gift from Cronan and Kuzminov, UIUC).

Transformants were selected on LB agar plates (10 g tryptone, 10 g NaCl, 5 g yeast extract per liter, plus 15 g agar) supplemented, where indicated, with kanamycin (5 *µ*g mL^−1^), chloramphenicol (10 *µ*g mL^−1^), both, or neither, according to the plasmid combination.

### Plate Reader Experiment

Overnight cultures were grown at 37 ^*◦*^C with 250 rpm orbital shaking in MOPS minimal medium supplemented with 1.12 mM K_2_HPO_4_, 0.2% (w/v) glucose, 0.1% (w/v) casamino acids, and thiamine (50 *µ*g mL^−1^). Where indicated, media contained kanamycin (5 *µ*g mL^−1^), chloramphenicol (10 *µ*g mL^−1^), both, or neither, matching the plasmid selection markers used in each experiment. Overnight cultures were diluted 1:100 into fresh supplemented MOPS and incubated for 3 h at 37 ^*◦*^C with the same shaking conditions.

Cells were then diluted with fresh MOPS to OD_600_ = 0.02 immediately before loading into the plate. A total of 200 *µ*L per condition was dispensed into a clear 96-well plate (catalog no. 3511724) with four biological replicates. Inducers were added at the indicated times: 3-oxo-C6-acyl-homoserine lactone (AHL; final 35 nM unless otherwise noted), anhydrotetracycline (aTc; 100 ng mL^−1^), and isopropyl *β*-D-1-thiogalactopyranoside (IPTG; 1 mM). OD_600_ and GFP fluorescence were measured every 5 min on a BioTek Synergy Neo2 plate reader maintained at 37 ^*◦*^C with continuous double-orbital shaking at 548 rpm between reads. The fluorescence intensity of GFP was measured using the internal monochromator in inverted (bottom-up) acquisition mode at an excitation wavelength of 473 nm, an emission wavelength of 520 nm, and a gain setting of 80.

### Analysis of Plate Reader Data

Plate-reader data (GFP and OD_600_) were exported as CSV files and analyzed in Python. GFP fluorescence was normalized by OD_600_, and background fluorescence was removed by subtracting the signal from empty-vector control wells that received the same inducers. For each condition (predefined well groups), biological replicates (*n* = 4) were averaged, and standard deviations were calculated. To quantify the degree to which the wave response of our various circuit designs resemble a square wave we leverage a mathematical generalization of the square wave, the trapezoidal pulse.

### Quantification of Experimentally Generated Wave Pulses

Trapezoidal and square waveform pulses are ubiquitous in modeling dynamics. In the context of dynamics as well as in representing fuzzy numbers, trapezoidal pulses generalize square waves. In the context of dynamic modeling trapezoidal pulses are considered a generalization of square pulses with potential for changing performance and energy efficiency. ^17^ In the context of representing fuzzy numbers, trapezoidal pulses are a fuzzy generalization of a square pulse, or “crisp interval”.^18^ Regardless of the application, trapezoidal pulses are five parameter generalizations of the three parameter square pulses. When the parameters of a trapezoidal pulse corresponding to ramp-up time and ramp-down time are zero it becomes a square pulse.

Each mean fluorescence trace 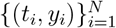 was fit to a such a five-parameter trapezoid pulse *y*(*t*; *θ*) with parameters *θ* = (*t*_0_, *t*_1_, *h, t*_2_, *t*_3_). The model represents a waveform that rises linearly from zero to height *h* between *t*_0_ and *t*_1_, remains constant at *h* from *t*_1_ to *t*_2_, and then decreases linearly back to zero between *t*_2_ and *t*_3_. Formally:

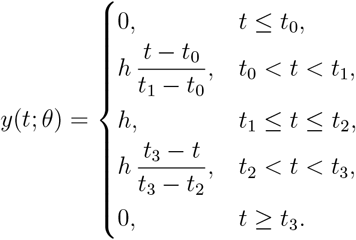

Parameters were obtained by constrained least squares, in which we minimize the objective function:

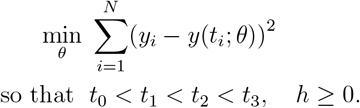

The primary timing metric, plateau duration, was reported as *τ*_plateau_ = *t*_2_ − *t*_1_.

### Single Cell Fluorescence Microscopy

Cells were revived from glycerol stock overnight at 37 ^*◦*^C in supplemented MOPS medium, diluted 1:100 into fresh MOPS, and recovered for 3 h to log phase. Cultures were then diluted to 5 × 10^6^ cells mL^−1^ in MOPS and loaded into a CellASIC plate. Separate solutions for flowing MOPS with 35 nM AHL and MOPS with 35 nM AHL and 100 ng mL^−1^ aTc were prepared and loaded into reagent wells in the CellASIC ONIX B04A plate for imaging. The plate was maintained at 37 ^*◦*^C during time-lapse microscopy. Images were cropped and preprocessed in Fiji, and single-cell fluorescence intensities were quantified in Python.

## Author Information

## Author Contributions

Jiayi Wu designed the genetic circuit, designed and conducted the experiments, analyzed the data, rendered figures, and wrote the manuscript. Charles Johnson assisted with the design of metrics for evaluating square wave-from dynamics and computation of metrics. Enoch Yeung assisted with the design of the research project, provided mentoring and feedback, and secured project funding. All authors approved the final version.

## Notes (Conflicts of Interest)

The authors declare no competing financial interest.

## Acknowledgment

The authors would like to thank Glen E. Cronan and Andrei Kuzminov for generously providing the E. coli Δ*sspB* strain and the SspB and GFP-DAS+4 plasmids used in this study, and express special thanks to Lili Yang and Yanran Wang for providing valuable feedback on experimental design, methods, and figure drafting.

