## Supplementary Information for "Tuning a Genetic Circuit with Double Negative Feedforward Loops to Approximate Square Waves"

### Supplementary Figures

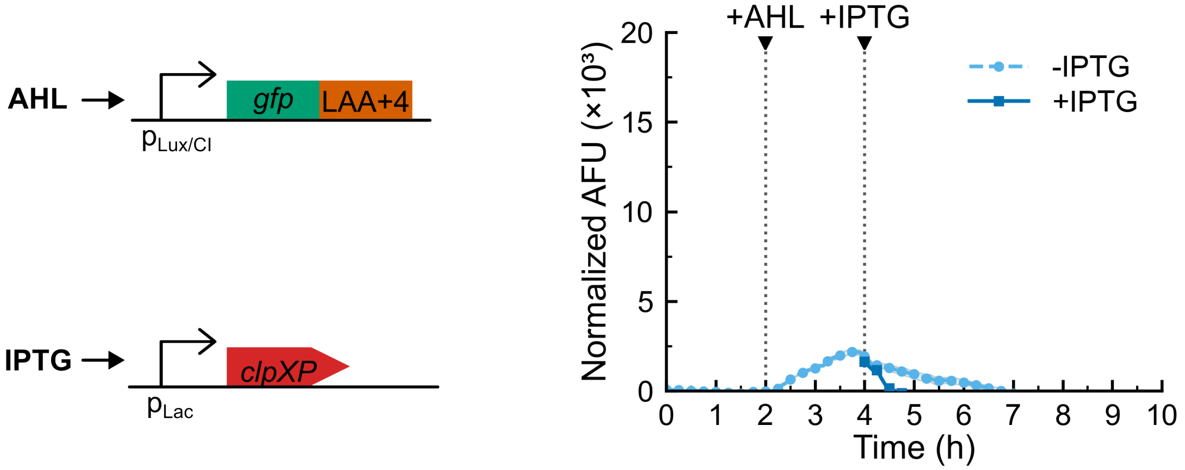


(a) (b)

Figure S1: **A high-affinity degron collapses dynamic range by increasing basal degradation. (a)** Circuit diagram for the LAA+4 control construct. LAA+4 substitutes the C-terminal DAS motif with an LAA motif, generating an adaptor-independent degron that is directly recognized by ClpXP. GFP–LAA+4 is expressed from the AHL-inducible $p_{\mathrm{Lux/CI}}$ promoter, and ClpXP is expressed from the IPTG-inducible $p_{\mathrm{Lac}}$ promoter. **(b)** Plate-reader time courses for the LAA+4 circuit. Normalized AFU trajectories from cultures induced with AHL (35 nM at 2 h) and, where indicated, IPTG (1 mM at 5 h). Data are background-subtracted and normalized; curves show mean ± SD ($n = 4$), with gray bands denoting SD. Vertical dotted lines mark inducer addition times; colors correspond to +IPTG and −IPTG conditions. Fluorescence remains low in both conditions, and IPTG further suppresses the already small pulse, indicating high basal degradation that collapses dynamic range and prevents accumulation of the pre-switch reservoir required for the sharp, high-amplitude switch observed with GFP–DAS+4 (Fig.5B).

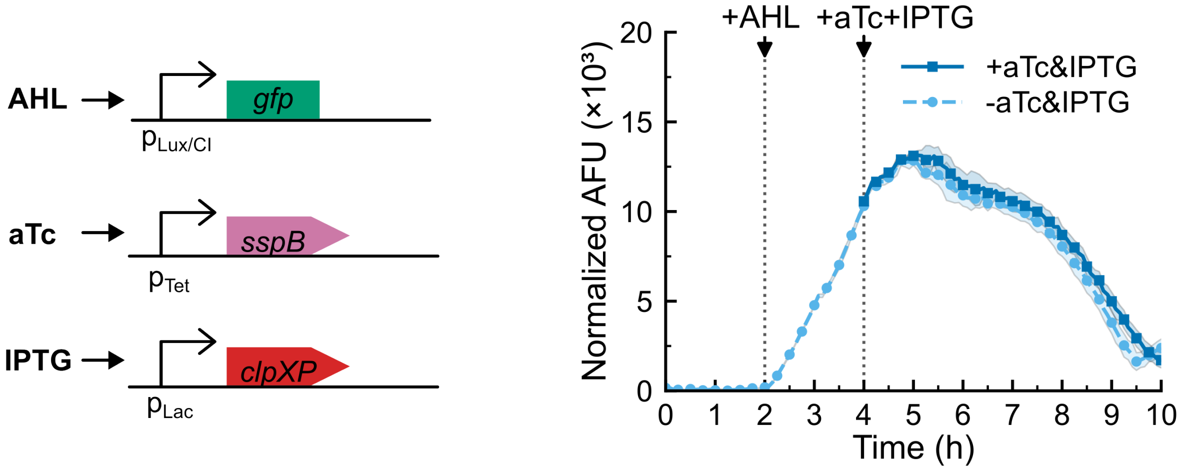


(a) (b)

Figure S2: **Orthogonality controls confirm degradation-pathway specificity. (a)** Schematic of the untagged GFP control construct. GFP lacking a degradation tag is expressed from the AHL-inducible $p_{\mathrm{Lux/CI}}$ promoter, while the degradation module components SspB and ClpXP are expressed from the aTc-inducible $p_{\mathrm{Tet}}$ and IPTG-inducible $p_{\mathrm{Lac}}$ promoters, respectively. **(b)**Plate-reader time courses for the untagged GFP control construct. Normalized AFU trajectories from cultures induced with AHL (35 nM at 2 h) and, where indicated, aTc (100 ng mL$^{-1}$) and IPTG (1 mM) at 4 h. Data are background-subtracted and normalized; curves show mean ± SD ($n = 4$), with gray bands denoting SD. Vertical dotted lines mark inducer addition times; colors correspond to +aTc/+IPTG and −aTc/−IPTG conditions. Co-induction of SspB and ClpXP produces no detectable change in amplitude, timing, or decay rate relative to the uninduced control, confirming that the accelerated OFF dynamics observed with GFP–DAS+4 (Fig.5B) require the DAS degron and do not arise from nonspecific cellular effects.

### Supplementary Material

#### 0.1 Sequence of the Reporter Plasmid Insert

**Legend for annotated reporter plasmid insert**

- prLacI promoter (prLacI)
- LuxR coding sequence (BBa_C0062)
- $p_{\mathrm{Lux/CI\text{-}OR}}$ promoter (BBa_K415032)
- GFP coding sequence
- Linker peptide sequence
- Myc epitope tag
- DAS+4 degron

```
CACCTCGAGTGGTGCAAAACCTTTCGCGGTATGGCATGATAGCGCCCGGAAGAGAGTCAA
TTCAGGGAAAGAGGAGAAATACTAGATGAAAAACATAAATGCCGACGACACATACAGAAT
AATTAATAAAATTAAAGCTTGTAGAAGCAATAATGATATTAATCAATGCTTATCTGATAT
GACTAAAATGGTACATTGTGAATATTATTTACTCGCGATCATTTATCCTCATTCTATGGT
TAAATCTGATATTTCAATCCTAGATAATTACCCTAAAAAATGGAGGCAATATTATGATGA
CGCTAATTTAATAAAATATGATCCTATAGTAGATTATTCTAACTCCAATCATTCACCAAT
TAATTGGAATATATTTGAAAACAATGCTGTAAATAAAAAATCTCCAAATGTAATTAAAGA
AGCGAAAACATCAGGTCTTATCACTGGGTTTAGTTTCCCTATTCATACGGCTAACAATGG
CTTCGGAATGCTTAGTTTTGCACATTCAGAAAAAGACAACTATATAGATAGTTTATTTTT
ACATGCGTGTATGAACATACCATTAATTGTTCCTTCTCTAGTTGATAATTATCGAAAAAT
AAATATAGCAAATAATAAATCAAACAACGATTTAACCAAAAGAGAAAAAGAATGTTTAGC
GTGGGCATGCGAAGGAAAAAGCTCTTGGGATATTTCAAAAATATTAGGTTGCAGTGAGCG
TACTGTCACTTTCCATTTAACCAATGCGCAAATGAAACTCAATACAACAAACCGCTGCCA
AAGTATTTCTAAAGCAATTTTAACAGGAGCAATTGATTGCCCATACTTTAAAAATTAATA
ACACTGATAGTGCTAGTGTAGATCACTACTAGAGAAAAAAAAACCCCGCCCCTGACAGGG
CGGGGTTTTTTTTGTCTCTTCAGTCCAATGGCGTGGACAATGCTTAATAAGCGATCCAGA
CGGGGCATGCAAATATTGGCTGGCGTATACTATCAATGTCTATTTCAGCCTATTCGTGCG
CGGTAATCTGTTGGTTGCGGTTGCGCGTATGCTTATTGTCTGTTATAAAGCGGCTATAAC
GCAAACCTTGATCTCCGGCCTTCTCACGCGTGTGGCTTAAACCTTTGGAGTTACGACACT
TGTCAGACTGGTACTAGATAATTGGATTGAGAGTGGTGCTCCACATTATGTTTCCACTCT
AGTCGCGGAGAGTACCTGTAGGATCGTACAGGTTTACGCAAGAAAATGGTTTGTTATAGT
CGAATACCTCTGGCGGTGATAAATTCATTAAAGAGGACAAAGGTACCATGCGTAAAGGAG
AAGAACTTTTCACTGGAGTTGTCCCAATTCTTGTTGAATTAGATGGTGATGTTAATGGGC
ACAAATTTTCTGTCAGTGGAGAGGGTGAAGGTGATGCAACATACGGAAAACTTACCCTTA
AATTTATTTGCACTACTGGAAAACTACCTGTTCCATGGCCAACACTTGTCACTACTTTCG
GTTATGGTGTTCAATGCTTTGCGAGATACCCAGATCATATGAAACAGCATGACTTTTTCA
AGAGTGCCATGCCCGAAGGTTATGTACAGGAAAGAACTATATTTTTCAAAGATGACGGGA
ACTACAAGACACGTGCTGAAGTCAAGTTTGAAGGTGATACCCTTGTTAATAGAATCGAGT
TAAAAGGTATTGATTTTAAAGAAGATGGAAACATTCTTGGACACAAATTGGAATACAACT
ATAACTCACACAATGTATACATCATGGCAGACAAACAAAAGAATGGAATCAAAGTTAACT
TCAAAATTAGACACAACATTGAAGATGGAAGCGTTCAACTAGCAGACCATTATCAACAAA
ATACTCCAATTGGCGATGGCCCTGTCCTTTTACCAGACAACCATTACCTGTCCACACAAT
```

CTGCCCTTTCGAAAGATCCCAACGAAAAGAGAGACCACATGGTCCTTCTTGAGTTTGTAA
CAGCTGCTGGGATTACACATGGCATGGATGAACTATACAAAAGTGCAGGTTCTGCAGCCG
GTTCTGGTGCAGCTGAACAAAAACTCATCTCTGAAGAAGATCTGGCCGCAAATGATGAAA
ACTACTCCGAGAATTATGCTGACGCGTCCTAATAAGCTTGATGGGGGATCCCATGGTACG
CGTGCTAGAGGCATCAAATAAAACGAAAGGCTCAGTCGAAAGACTGGGCCTTTCGTTTTA
TCTGTTGTTTGTCGGTGAACGCTCTCCTGAGTAGGACAAAT

#### 0.2 Sequence of the Regulatory Plasmid Insert

##### Legend for annotated regulatory plasmid insert

- Constitutive promoter pJ23150
- lacI coding sequence
- TetR coding sequence
- pTet promoter
- sspB coding sequence
- mf-Lon degradation tag (pdt)
- λCI coding sequence
- ClpX coding sequence
- ClpP coding sequence

GCTGTTCTTTACGGCTAGCTCAGTCCTAGGTATTATGCAAGCGGGCCCAAGTTCACTTAA
AAAGGAGATCAACAATGAAAGCAATTTTCGTACTGAAACATCTTAATCATGCACAGGAGA
CTTTCTAATGAAACCAGTAACGTTATACGATGTCGCAGAGTATGCCGGTGTCTCTTATCA
GACCGTTTCCCGCGTGGTGAACCAGGCCAGCCACGTTTCTGCGAAAACGCGGGAAAAAGT
GGAAGCGGCGATGGCGGAGCTGAATTACATTCCCAACCGCGTGGCACAACAACTGGCGGG
CAAACAGTCGTTGCTGATTGGCGTTGCCACCTCCAGTCTGGCCCTGCACGCGCCGTCGCA
AATTGTCGCGGCGATTAAATCTCGCGCCGATCAACTGGGTGCCAGCGTGGTGGTGTCGAT
GGTAGAACGAAGCGGCGTCGAAGCCTGTAAAGCGGCGGTGCACAATCTTCTCGCGCAACG
CGTCAGTGGGCTGATCATTAACTATCCGCTGGATGACCAGGATGCCATTGCTGTGGAAGC
TGCCTGCACTAATGTTCCGGCGTTATTTCTTGATGTCTCTGACCAGACACCCATCAACAG
TATTATTTTCTCCCATGAGGACGGTACGCGACTGGGCGTGGAGCATCTGGTCGCATTGGG
TCACCAGCAAATCGCGCTGTTAGCGGGCCCATTAAGTTCTGTCTCGGCGCGTCTGCGTCT
GGCTGGCTGGCATAAATATCTCACTCGCAATCAAATTCAGCCGATAGCGGAACGGGAAGG
CGACTGGAGTGCCATGTCCGGTTTTCAACAAACCATGCAAATGCTGAATGAGGGCATCGT
TCCCACTGCGATGCTGGTTGCCAACGATCAGATGGCGCTGGGCGCAATGCGCGCCATTAC
CGAGTCCGGGCTGCGCGTTGGTGCGGATATCTCGGTAGTGGGATACGACGATACCGAGGA
CAGCTCATGTTATATCCCGCCGTTAACCACCATCAAACAGGATTTTCGCCTGCTGGGGCA
AACCAGCGTGGACCGCTTGCTGCAACTCTCTCAGGGCCAGGCGGTGAAGGGCAATCAGCT
GTTGCCCGTCTCACTGGTGAAAAGAAAAACCACCCTGGCGCCCAATACGCAAACCGCCTC
TCCCCGCGCGTTGGCCGATTCATTAATGCAGCTGGCACGACAGGTTTCCCGACTGGAAAG
CGGGCAGTGAACTAACGCATGAGAAAGCCCCCGGAAGATCACCTTCCGGGGGCTTTTTTA
TTGCGCACGTCTCATTTTCGCCAGATATCGACGTCTTAAGACCCACTTTCACATTTAAGT
TGTTTTTCTAATCCGCATATGATCAATTCAAGGCCGAATAAGAAGGCTGGCTCTGCACCT

TGGTGATCAAATAATTCGATAGCTTGTCGTAATAATGGCGGCATACTATCAGTAGTAGGT
GTTTCCCTTTCTTCTTTAGCGACTTGATGCTCTTGATCTTCCAATACGCAACCTAAAGTA
AAATGCCCCACAGCGCTGAGTGCATATAATGCATTCTCTAGTGAAAAACCTTGTTGGCAT
AAAAAGGCTAATTGATTTTCGAGAGTTTCATACTGTTTTTCTGTAGGCCGTGTACCTAAA
TGTACTTTTGCTCCATCGCGATGACTTAGTAAAGCACATCTAAAACTTTTAGCGTTATTA
CGTAAAAAATCTTGCCAGCTTTCCCCTTCTAAAGGGCAAAAGTGAGTATGGTGCCTATCT
AACATCTCAATGGCTAAGGCGTCGAGCAAAGCCCGCTTATTTTTTACATGCCAATACAAT
GTAGGCTGCTCTACACCTAGCTTCTGGGCGAGTTTACGGGTTGTTAAACCTTCGATTCCG
ACCTCATTAAGCAGCTCTAATGCGCTGTTAATCACTTTACTTTTATCTAATCTAGACATA
CATTAATTCCTAATTTTTGTTGACACTCTATCGTTGATAGAGTTATTTTACCACTCCCTA
TCAGTGATAGAGAAAAGAATTCAAAAGATCTAAAGAGGAGAAAGGATCTATGGATTTGTC
ACAGCTAACACCACGTCGTCCCTATCTGCTGCGTGCATTCTATGAGTGGTTGCTGGATAA
CCAGCTCACGCCGCACCTGGTGGTGGATGTGACGCTCCCTGGCGTGCAGGTTCCTATGGA
ATATGCGCGTGACGGGCAAATCGTACTCAACATTGCGCCGCGTGCTGTCGGCAATCTGGA
ACTGGCGAATGATGAGGTGCGCTTTAACGCGCGCTTTGGTGGCATTCCGCGTCAGGTTTC
TGTGCCGCTGGCTGCCGTGCTGGCTATCTACGCCCGTGAAAATGGCGCAGGCACGATGTT
TGAGCCTGAAGCTGCCTACGATGAAGATACCAGCATCATGAATGATGAAGAGGCATCGGC
AGACAACGAAACCGTTATGTCGGTTATTGATGGCGACAAGCCAGATCACGATGATGACAC
TCATCCTGACGATGAACCTCCGCAGCCACCACGCGGTGGTCGACCGGCATTACGCGTTGT
GAAGGCGGCGAACAAAAACGAAGAAAACACCAACGAAGTGCCGACCTTTATGCTGAACGC
GGGCCAGGCGAACTATGCGTTTGCGTAATAAATTAAAGAGGAGAAAATGAGCACAAAAAA
GAAACCATTAACACAAGAGCAGCTTGAGGACGCACGTCGCCTTAAAGCAATTTATGAAAA
AAAGAAAAATGAACTTGGCTTATCCCAGGAATCTGTCGCAGACAAGATGGGGATGGGGCA
GTCAGGCGTTGGTGCTTTATTTAATGGCATCAATGCATTAAATGCTTATAACGCCGCATT
GCTTGCAAAAATTCTCAAAGTTAGCGTTGAAGAATTTAGCCCTTCAATCGCCAGAGAAAT
CTACGAGATGTATGAAGCGGTTAGTATGCAGCCGTCACTTAGAAGTGAGTATGAGTACCC
TGTTTTTTCTCATGTTCAGGCAGGGATGTTCTCACCTGAGCTTAGAACCTTTACCAAAGG
TGATGCGGAGAGATGGGTAAGCACAACCAAAAAAGCCAGTGATTCTGCATTCTGGCTTGA
GGTTGAAGGTAATTCCATGACCGCACCAACAGGCTCCAAGCCAAGCTTTCCTGACGGAAT
GTTAATTCTCGTTGACCCTGAGCAGGCTGTTGAGCCAGGTGATTTCTGCATAGCCAGACTT
GGGGGTGATGAGTTTACCTTCAAGAAACTGATCAGGGATAGCGGTCAGGTGTTTTTACAA
CCACTAAACCCACAGTACCCAATGATCCCATGCAATGAGAGTTGTTCCGTTGTGGGGAAA
GTTATCGCTAGTCAGTGGCCTGAAGAGACGTTTGGCTGAGGTACCTCACACTGGCTCACC
TTCGGGTGGGCCTTTCTGCGTTTATATACTAGAGAGAGAATATAAAAAGCCAGATTATTA
ATCCGGCTTTTTTATTATTTATAAATGTGAGCGGATAACATTGACATTGTGAGCGGATAA
CAAGATACTGAGCACATCAGCAGGACGCACTGACCGGGCCCAAGTTCACTTAAAAAGGAG
ATCAACAATGAAAGCAATTTTCGTACTGAAACATCTTAATCATGCTGAGGAAAGTTTCTA
ATGTCATACAGCGGCGAACGAGATAACTTTGCACCCCATATGGCGCTGGTGCCGATGGTC
ATTGAACAGACCTCACGCGGTGAGCGCTCTTTTGATATCTATTCTCGTCTACTTAAGGAA
CGCGTCATTTTTCTGACTGGCCAGGTTGAAGACCACATGGCTAACCTGATTGTGGCGCAG
ATGCTGTTCCTGGAAGCGGAAAACCCAGAAAAAGATATCTATCTGTACATTAACTCCCCA
GGCGGGGTGATCACTGCCGGGATGTCTATCTATGACACCATGCAGTTTATCAAGCCTGAT
GTCAGCACCATCTGTATGGGCCAGGCGGCCTCGATGGGCGCTTTCTTGCTGACCGCAGGG
GCAAAAGGTAAACGTTTTTGCCTGCCGAATTCGCGCGTGATGATTCACCAACCGTTGGGC
GGCTACCAGGGACAGGCGACCGATATCGAAATTCATGCCCGTGAAATTCTGAAAGTTAAA

GGGCGCATGAATGAACTTATGGCGCTTCATACGGGTCAATCATTAGAACAGATTGAACGT
GATACCGAGCGCGATCGCTTCCTTTCCGCCCCTGAAGCGGTGGAATACGGTCTGGTCGAT
TCGATTCTGACCCATCGTAATTGATAATACTAGATGCATTAATAATCACACTGGCTCACC
TCTCTACTGTTTCTCCATAGGGCCCAAGTTCACTTAAAAAGGAGATCAACAATGAAAGCA
ATTTTCGTACTGAAACATCTTAATCATGCGGAGGAGGGTTTCTAATGACAGATAAACGCA
AAGATGGCTCAGGCAAATTGCTGTATTGCTCTTTTTGCGGCAAAAGCCAGCATGAAGTGC
GCAAGCTGATTGCCGGTCCATCCGTGTATATCTGCGACGAATGTGTTGATTTATGTAACG
ACATCATTCGCGAAGAGATTAAAGAAGTTGCACCGCATCGTGAACGCAGTGCGCTACCGA
CGCCGCATGAAATTCGCAACCACCTGGACGATTACGTTATCGGCCAGGAACAGGCGAAAA
AAGTGCTGGCGGTCGCGGTATACAACCATTACAAACGTCTGCGCAACGGCGATACCAGCA
ATGGCGTCGAGTTGGGCAAAAGTAACATTCTGCTGATCGGTCCGACCGGTTCCGGTAAAA
CGCTGCTGGCTGAAACGCTGGCGCGCCTGCTGGATGTTCCGTTCACCATGGCCGACGCGA
CTACACTGACCGAAGCCGGTTATGTGGGTGAAGACGTTGAAAACATCATTCAGAAGCTGT
TGCAGAAATGCGACTACGATGTCCAGAAAGCACAGCGTGGTATTGTCTACATCGATGAAA
TCGACAAGATTTCTCGTAAGTCAGACAACCCGTCCATTACCCGAGACGTTTCCGGTGAAG
GCGTACAGCAGGCACTGTTGAAACTGATCGAAGGTACGGTAGCTGCTGTTCCACCGCAAG
GTGGGCGTAAACATCCGCAGCAGGAATTCTTGCAGGTTGATACCTCTAAGATCCTGTTTA
TTTGTGGCGGTGCGTTTGCCGGTCTGGATAAAGTGATTTCCCACCGTGTAGAAACCGGCT
CCGGCATTGGTTTTGGCGCGACGGTAAAAGCGAAGTCCGACAAAGCAAGCGAAGGCGAGC
TGCTGGCGCAGGTTGAACCGGAAGATCTGATCAAGTTTGGTCTTATCCCTGAGTTTATTG
GTCGTCTGCCGGTTGTCGCAACGTTGAATGAACTGAGCGAAGAAGCTCTGATTCAGATCC
TCAAAGAGCCGAAAAACGCCCTGACCAAGCAGTATCAGGCGCTGTTTAATCTGGAAGGCG
TGGATCTGGAATTCCGTGACGAGGCGCTGGATGCTATCGCTAAGAAAGCGATGGCGCGTA
AAACCGGTGCCCGTGGCCTGCGTTCCATCGTAGAAGCCGCACTGCTCGATACCATGTACG
ATCTGCCGTCCATGGAAGACGTCGAAAAAGTGGTTATCGACGAGTCGGTAATTGATGGTC
AAAGCAAACCGTTGCTGATTTATGGCAAGCCGGAAGCGCAACAGGCATCTGGTGAATAAA
AAACGGCGGGGTGATTGCCCCGCCGTTTTTTAGTGATGTGATGAGATGTGCAGCTTCTTT
TTTCGATACATGAGCGGAGTCATCTTCATTTCTTTTCGAAACTGATTAATAAAATAGCTT
GTACTATTAAACCCTACTTGATAGGCGACTTCCGTCACATTTGCTTCTTCTTGCTGCAGG
ATCCGGCTGCTAACAAAGCCCGAAAGGAAGCTGAGTTGGCTGCTGCCACCGCTGAGCAAT
AACTAGCATAACCCCTTGGGGCCTCTAAACGGGTCTTGAGGGGTTTTTTG
